# Comparative Biochemical and Structural Analysis of Novel Cellulose Binding Proteins (Tāpirins) from Extremely Thermophilic *Caldicellulosiruptor* Species

**DOI:** 10.1101/359786

**Authors:** Laura L. Lee, William S. Hart, Vladimir V. Lunin, Markus Alahuhta, Yannick J. Bomble, Michael E. Himmel, Sara E. Blumer-Schuette, Michael W.W. Adams, Robert M. Kelly

**Affiliations:** Department of Chemical and Biomolecular Engineering North Carolina State University, Raleigh, NC 27695; Biosciences Center, National Renewable Energy Laboratory, Golden, CO, 80401; Department of Biochemistry and Molecular Biology University of Georgia, Athens, Georgia 30602

**Keywords:** Caldicellulosiruptor, tāpirins, lignocellulose, glycoside hydrolase, cellulase

## Abstract

Genomes of extremely thermophilic *Caldicellulosiruptor* species encode novel cellulose binding proteins, tāpirins, located proximate to the type IV pilus locus. Previously, the *C*-terminal domain of a tāpirin (Calkro_0844) from *Caldicellulosiruptor kronotskyensis* was shown to be structurally unique and have a cellulose binding affinity akin to family 3 carbohydrate binding modules (CBM3). Here, full-length and C-terminal versions of tāpirins from *Caldicellulosiruptor bescii* (Athe_1870), *Caldicellulosiruptor hydrothermalis* (Calhy_0908), *Caldicellulosiruptor kristjanssonii* (Calkr_0826), and *Caldicellulosiruptor naganoensis* (NA10_0869) were produced recombinantly in *Escherichia coli* and compared to Calkro_0844. All five tāpirins bound to microcrystalline cellulose, switchgrass, poplar, filter paper, but not to xylan. Densitometry analysis of bound protein fractions visualized by SDS-PAGE revealed that Calhy_0908 and Calkr_0826 (from weakly cellulolytic species) associated with the cellulose substrates to a greater extent than Athe_1870, Calkro_0844 and NA10_0869 (from strongly cellulolytic species), perhaps to associate closely with biomass to capture glucans released from lignocellulose by cellulases produced in *Caldicellulosiruptor* communities. Three-dimensional structures of the C-terminal binding regions of Calhy_0908 and Calkr_0826 were closely related to Calkro_0844, despite the fact that their amino acid sequence identities compared to Calkro_0844 were only 16% and 36%, respectively. Unlike the parent strain, *C. bescii* mutants lacking the tāpirin genes did not bind to cellulose following short-term incubation, reinforcing the significance of these proteins in cell association with plant biomass. Given the scarcity of carbohydrates in neutral terrestrial hot springs, tāpirins likely help cells scavenge carbohydrates from lignocellulose to support growth and survival of *Caldicellulosiruptor* species.

**Importance:** Mechanisms by which microorganisms attach to and degrade lignocellulose are important to understand if effective approaches for conversion of plant biomass into fuels and chemicals are to be developed. *Caldicellulosiruptor* species grow on carbohydrates from lignocellulose at elevated temperatures and have biotechnological significance for that reason. Novel cellulose binding proteins, called tāpirins, are involved in the way *Caldicellulosiruptor* species interact with microcrystalline cellulose and here additional information about the diversity of these proteins across the genus is provided, including three dimensional structural comparisons.

## Introduction

The natural capacity to utilize both the cellulose and hemicellulose content of plant biomass as microbial growth substrates is relatively rare, especially among extreme thermophiles growing optimally above 70°C (1). However, in pH neutral, terrestrial hot springs and thermal features, species from the genus *Caldicellulosiruptor* can be isolated, all of which utilize hemicellulose, but only some of which can hydrolyze microcrystalline cellulose (2, 3). To degrade plant material, *Caldicellulosiruptor* species draw from an inventory of intracellular, surface (S)-layer associated, and secreted glycoside hydrolases (GHs) with complementary modes of action (4, 5). Unlike cellulosomal or free enzyme systems commonly found in other cellulolytic organisms, such as *Clostridia* (6) and *Trichoderma* (7), respectively, many *Caldicellulosiruptor* carbohydrate active enzymes (CAZymes (8)) are modular and consist of combinations of catalytic and non-catalytic (e.g., carbohydrate binding module [CBM]) domains connected by proline/threonine-rich linkers in various arrangements. The most well studied example is the multi-functional cellulase, CelA, which is arranged as GH9-CBM3-CBM3-CBM3-GH48 domains, where the numbers refer to specific protein families (9-14). The synergy between the endoglucanase (GH9) and exoglucanase (GH48) domains contributes to a novel mode of action for CelA that involves physical burrowing into cellulose fibers, thereby creating cavities for further enzymatic access to the carbohydrate content of plant biomass (9).

While not directly responsible for lignocellulose degradation, the non-catalytic domains in CelA also play an important role. In general, CBMs improve the efficacy of GHs by ensuring proximity to the substrate, as well as in some cases contributing to thermostability (15, 16). CBM3s, in particular, are specific to cellulose and allow enzymes like CelA to attach to their substrates such that their GH domains are proximate to their substrate (9). Other non-catalytic protein features in the *Caldicellulosiruptor* also play a role in orienting cells to lignocellulosic carbohydrates. S-layer homology (SLH) domains are associated with certain modular GHs in these bacteria (17, 18). For instance, Calkro_0402, a xylanase with GH10, CBM22, and CBM9 domains, is anchored to the cell surface of the strongly cellulolytic *Caldicellulosiruptor kronotskyensis* and the gene encoding this enzyme is highly transcribed during growth on lignocellulose (switchgrass). When inserted into the genome of *Caldicellulosiruptor bescii*, Calkro_0402 improved the attachment of cells to xylan and significantly increased xylan degradation, despite the fact that wild type *C. bescii* produces other xylanases (17).

In addition to CBMs within modular enzymes, *Caldicellulosiruptor* species also use non-catalytic proteins to bind to lignocellulosic substrates. Transcriptomic and proteomic analysis of cellulose-bound *Caldicellulosiruptor* cultures identified the presence of carbohydrate binding proteins (19), including the recently characterized ‘tāpirins’ (20). Tāpirins typically have a Mr of approximately 70 kDa as a single polypeptide; recombinant versions from *C. kronotskyensis* have a specific affinity for cellulose fibers in plant material and an affinity for Avicel similar to that of CBM3s, despite no significant structural homology (20). While the full-length structure has not been resolved, the cellulose-binding, 38.4 kDa C-terminal domain from Calkro_0844 was successfully crystallized and shown to be novel within the current protein database. Hydrophobic and aromatic residues present on the face of a β-helix likely make up the binding pocket with a flexible loop overhanging, and potentially protecting, access to it (20). Interestingly, tāpirin genes are located near the type IV pilus (T4P) locus in *Caldicellulosiruptor* species, suggesting a potential functional connection.

In the present study, comparative assessment of tāpirins across the genus *Caldicellulosiruptor* was conducted (see **Figure 1**). Structural data for two additional tāpirins from less cellulolytic species are provided, as is an assessment of relative binding capacities. Additionally, the role of the tāpirins was further explored through gene deletions in *C. bescii* and the resulting impact of binding.

**Figure 1.**
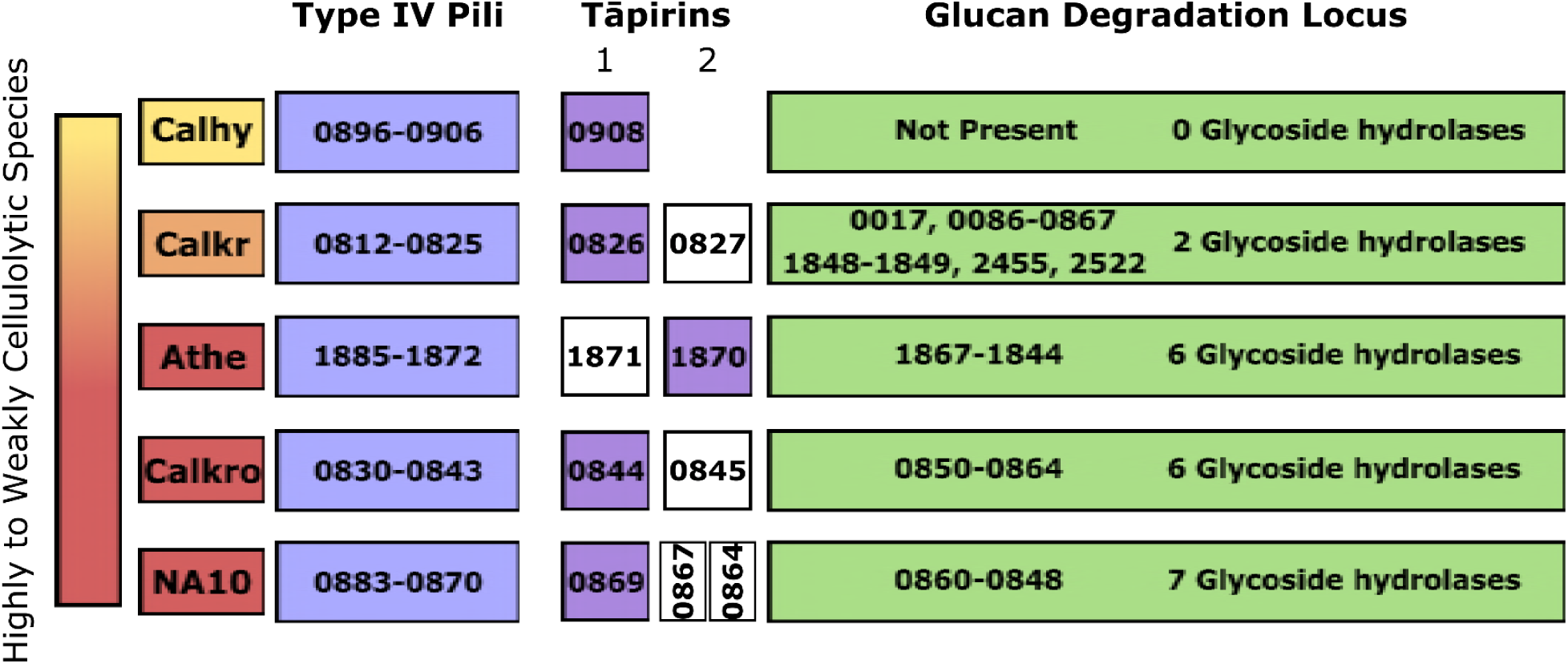
Genomic organization of Type IV pili, tāpirins, and glucan degradation loci in examined *Caldicellulosiruptor* species. Numbers refer to gene locus tags in sectioned loci. The tāpirins that were examined in this study are highlighted in purple. Abbreviations are as follows: Athe, *C. bescii*; Calhy, *C. hydrothermalis*; Calkr, *C. kristjanssonii*; Calkro, *C. kronotskyensis*; NA10, *C. naganoensis*.

## Results and Discussion

### Tapirins and pili are necessary for rapid binding to cellulosic substrates

Degradation of lignocellulosic substrates by *Caldicellulosiruptor* populations is likely predicated on substrate attachment, as genes encoding for key cellulases from the Glucan Degradation Locus (GDL) (21) are genomic neighbors of a T4P locus and tāpirins **(Figure 1)** (20). To assess the importance of the tāpirins and/or T4P in binding to cellulose, knockouts of tāpirins (Athe_1870-1871) and the entire pilus locus plus tāpirins (Athe_1870-1885) were generated in *C. bescii.* After an hour incubation with Avicel, unlike the parent strain, both knockouts showed no propensity for binding, based on changes on planktonic cell densities **(Figure 2).** Statistically insignificant changes were noted in the planktonic cell density for the tapirins and tapirins/T4P KOs when grown in the presence of microcrystalline cellulose, while in the parent strain planktonic cell density was reduced by more than 7 x 10^7^ cells/ml. This suggests that the tāpirins and T4P play a role in cell adherence to cellulosic substrates, at least during initial exposure, which could be critical for scavenging carbohydrates in otherwise nutrient-limited hot springs.

**Figure 2.**
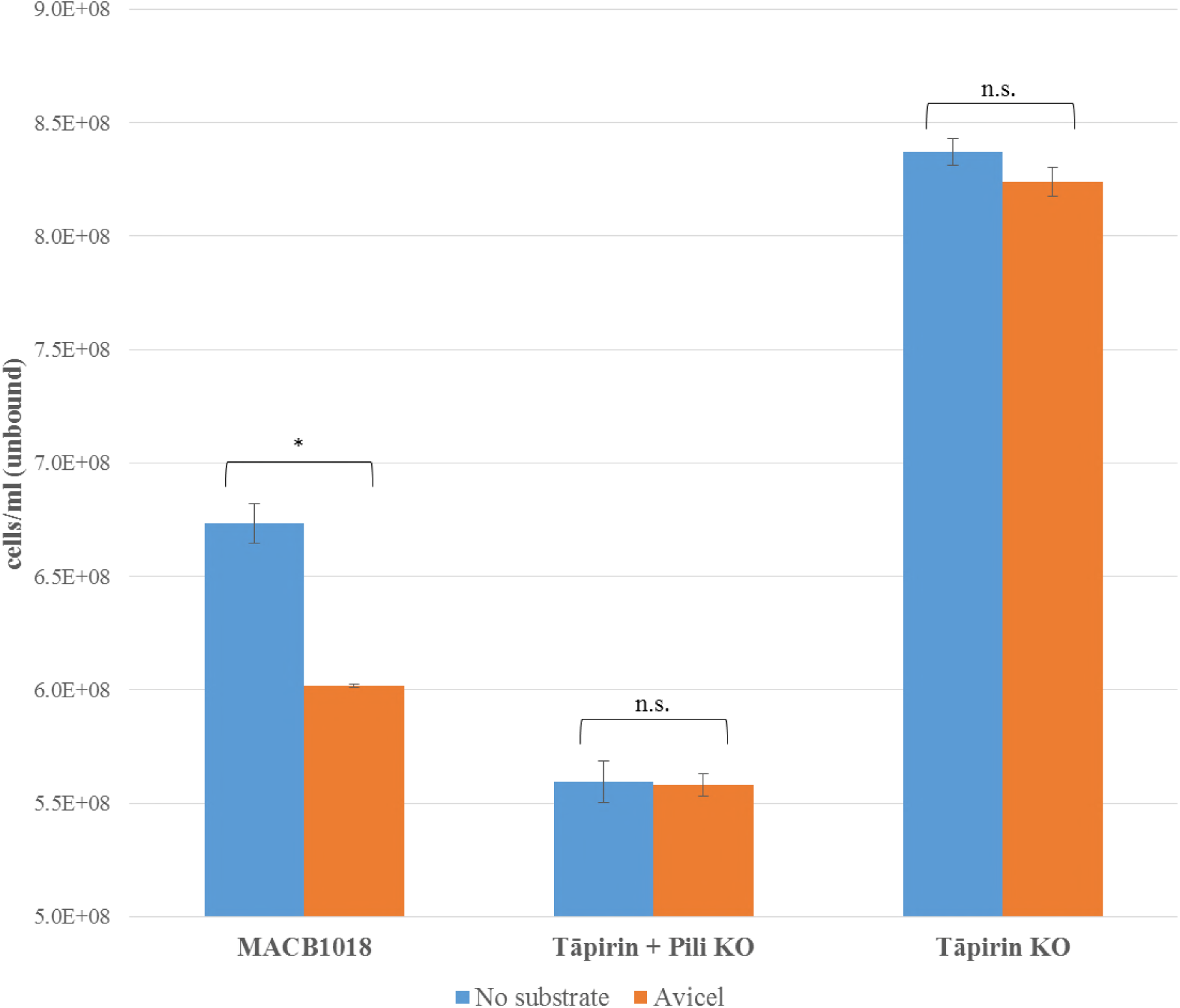
Whole cell binding assay of parent *Caldicellulosiruptor bescii* strain, MACB1018, versus *C. bescii* tāpirin and pili deletion (ΔAthe_1870-1885 - RKCB135) and tāpirin only deletion (ΔAthe_1870-1871 - RKCB136) strains. Cell concentrations refer to cells not attached to Avicel and/or test tube wall after 1 h incubation. Error bars are standard error of triplicate samples (* indicates statistical significance; ‘n.s.’ is ‘not significant’).

### Tāpirins are ubiquitous in the genus *Caldicellulosiruptor*

Previous studies showed that putative tāpirins are found in all genome-sequenced *Caldicellulosiruptor* species and were assigned to one of two groups (Class 1 and Class 2), based on their localization with genes encoding; class 1 tāpirins are closest to the T4P locus and class 2 tāpirins (if there are more than one tāpirin) closest to the GDL **(Figure 1)** (20). The GDL encodes up to seven glycoside hydrolases (GHs), which collectively contain GH5, GH9, GH10, GH12. GH44, GH48, GH74, and CBM3 domains, and these enzymes plays an essential role in microcrystalline hydrolysis (21). The most cellulolytic *Caldicellulosiruptor* species (e.g., *C. bescii, C. kronotskyensis*, and *C. naganoensis)* produce six to seven GDL GHs, while the less cellulolytic species produce an incomplete set of these enzymes. For example, *C. kristjanssonii*, which is less cellulolytic, produces two of the GDL enzymes, while *C. hydrothermalis*, which minimally degrades microcrystalline cellulose, lacks all of the GDL GHs **(Figure 1).** Amino acid sequence analysis indicates that while some tāpirins are closely related (e.g., Calkro_0844 and NA10_0869 from two highly cellulolytic species, *C. kronotskyensis* and *C. naganoensis*, are 85% identical at the amino acid level), Calhy_0908 from the weakly cellulolytic *C. hydrothermalis* is less than 18% identical to the other tāpirins. Interestingly, the N-termini of these tāpirins appear to share the most homology across the whole protein sequence (seen with residues 1-298 in **Figure 3,** on the right side of the indicated linker), with the exception of Calhy_0908; in fact, Athe_1870 shares 86% similarity in this N-terminal range with both Calkro_0844 and NA10_0869 **(Table 2A).** Overall, the N-terminus of the tāpirin may be responsible for how tāpirins, in general, associate with the cell surface, while the C-terminus (%ID in **Table 2B**) establishes the binding function, the latter of which was determined to be the case for Calkro_0844 (20).

**Figure 3.**
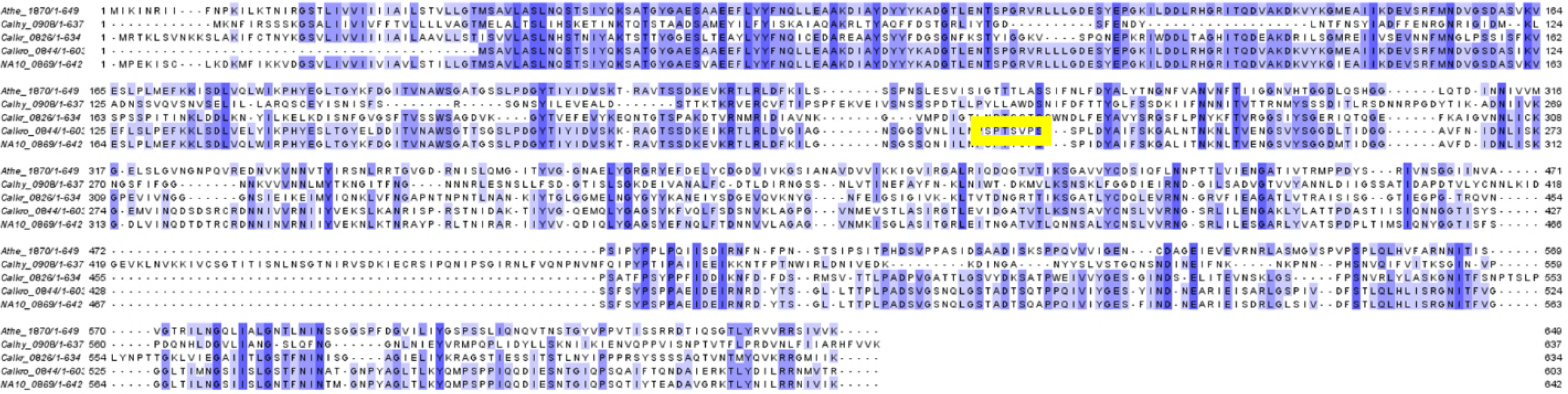
Pileup of full length amino acid sequences of tāpirins Athe_1870, Calhy_0908, Calkr_0826, Calkro_0844, and NA10_0869. Calhy_0908, Calkr_0826, and Calkro_0844 were aligned based on structural homology, while Athe_1870 and NA10_0869 were aligned with NCBI-BLAST (25). The darker the color purple is, the more shared homology there is between those specific sequences (actual amino acid %ID between tāpirins are listed in **Tables 1** and **2A-B**. The yellow box represents the location of the linker (residues 269-275 in alignment, and residues 257-264 in full length Calkro_0844 sequence) between the N- and C-terminal of Calkro_0844.

**Table 1.**
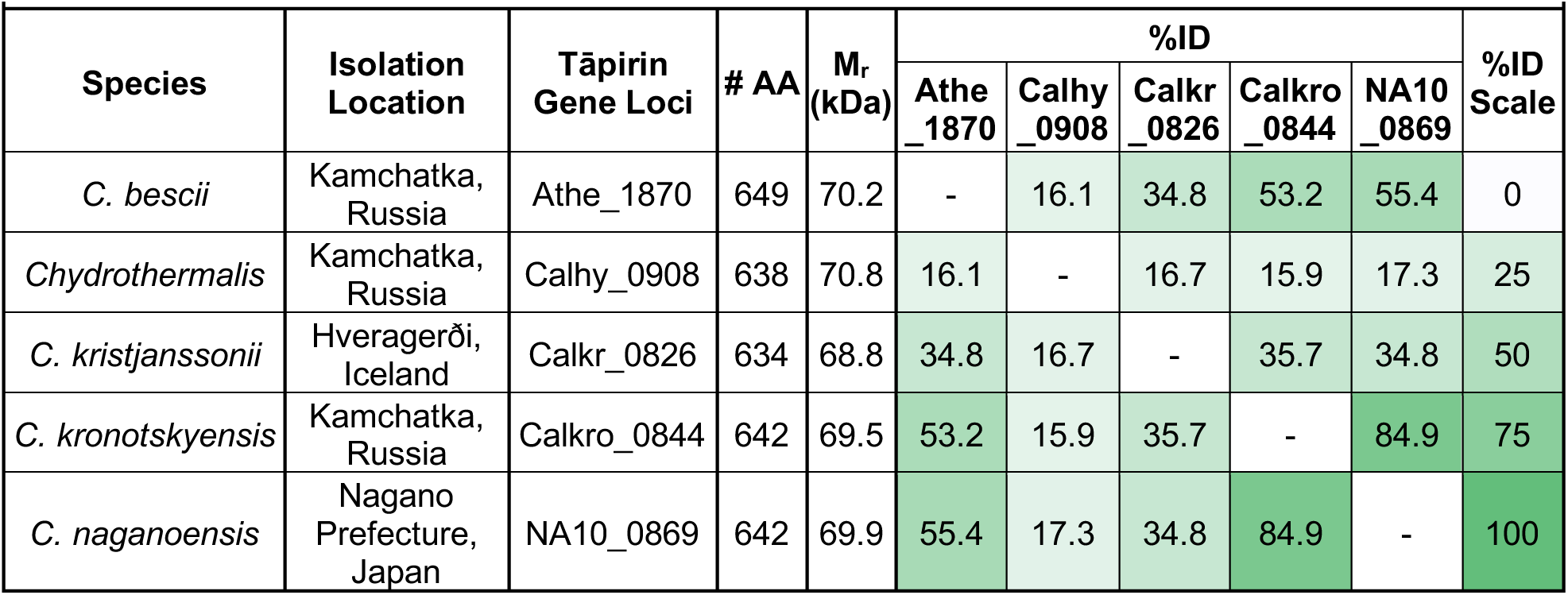
Characteristics of selected tāpirins from *Caldicellulosiruptor* species

**Table 2:**
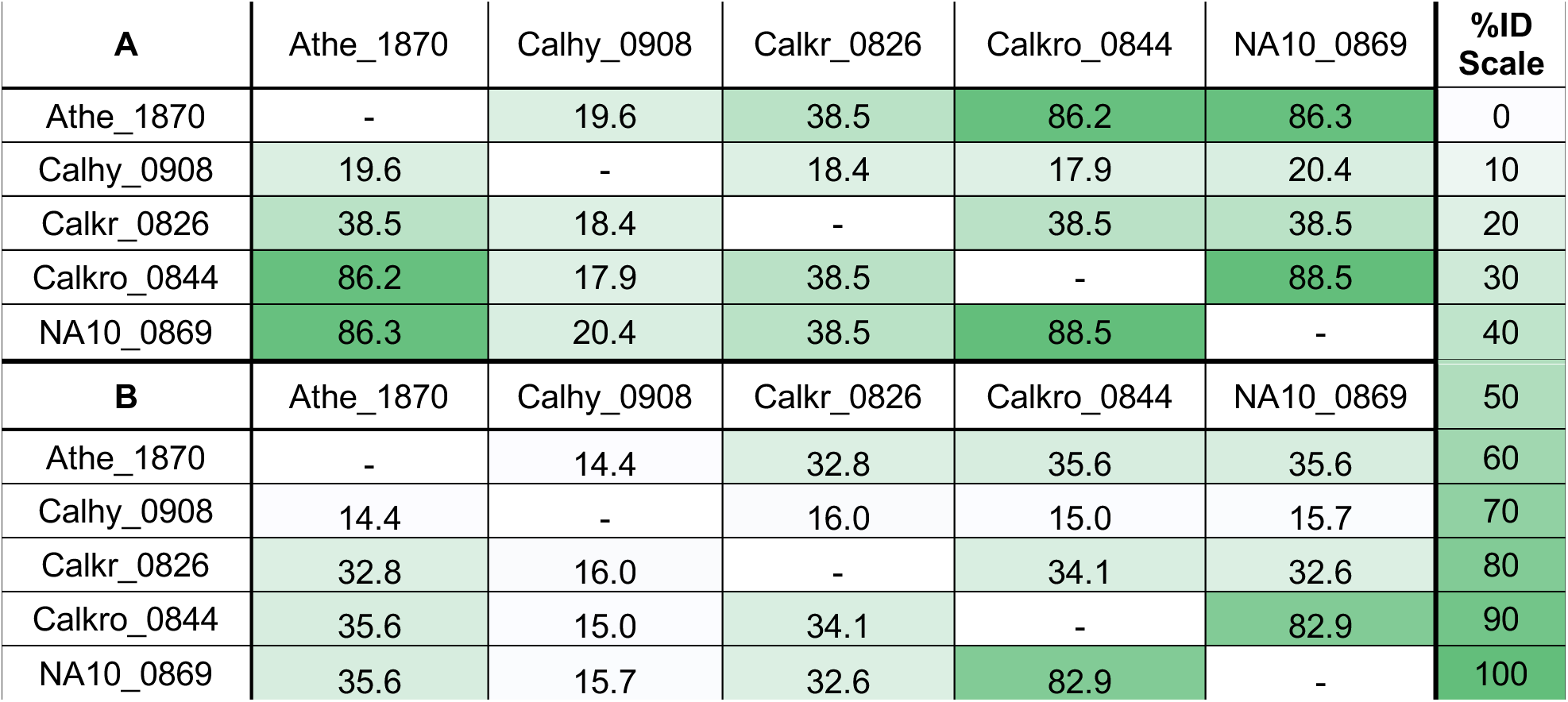
Sequence ID across the A) *N*-termini and B) *C*-termini of Athe_1870, Calhy_0908, Calkr_0826, Calkro_0844, and NA10_0869. Residues 1-268 (A) and 276-758 (B) from the full length alignment (**Figure 3**) were used in these comparisons. Athe_1870, Calhy_0908, Calkr_0826, Calkro_0844, and NA10_0869 are from *C. bescii*, *C. hydrothermalis*, *C. kristjanssonii*, *C. kronotskyensis*, and *C. naganoensis*, respectively.

### *In vitro* binding assays with tāpirins and plant-based substrates

Tāpirins were initially characterized from *C. kronotskyensis* (Calkro_0844 and Calkro_0845) and from *Caldicellulosiruptor saccharolyticus* (Csac_1073) (20). Binding assays showed that tāpirins from these two species preferentially adhered to cellulose. To confirm that this substrate specificity was consistent across the genus *Caldicellulosiruptor*, four additional tāpirins from other species were recombinantly produced to examine their binding to plant biomass-related substrates (see **Figure 4).** Athe_1870, Calhy_0809, Calkr_0826, and NA10_0869 from *C. bescii, C. hydrothermalis, C. kristjanssonii*, and *C. naganoensis*, respectively, along with the previously characterized Calkro_0844, were incubated with Avicel, filter paper and xylan, as well as with lignocellulose (switchgrass and poplar). After incubation, the samples were split into ‘unbound’ and ‘substrate-bound’ fractions, both of which were visualized with SDS-PAGE. No tāpirins bound to xylan to any significant extent, but consistently adhered to the other substrates. It was interesting that Calkr_0826 and Calhy_0908 bound to poplar and switchgrass to a greater extent than the tāpirins from more strongly cellulolytic species. In fact, the tāpirin from the least cellulolytic species tested, Calhy_0908, appears to adhere to more binding sites on cellulosic materials overall.

**Figure 4.**
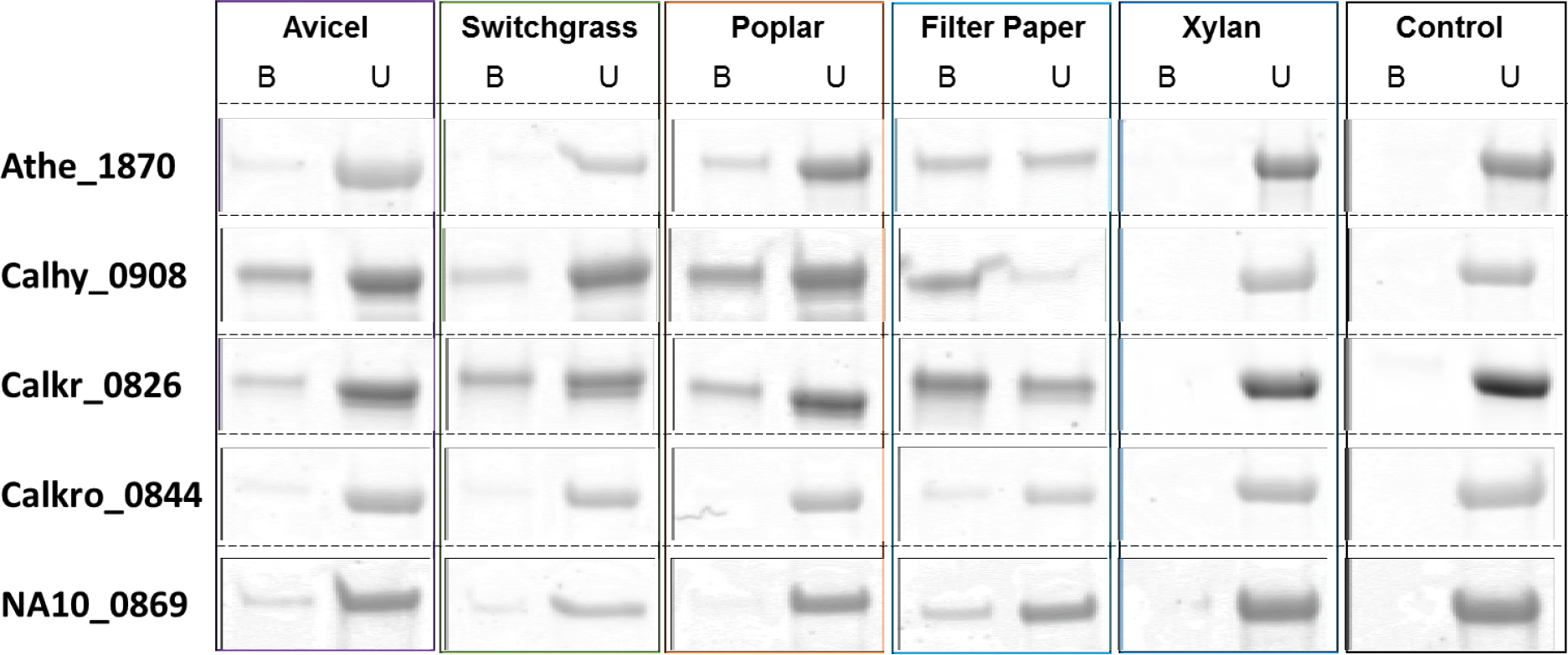
SDS-PAGE gel of tāpirin binding assay to plant component substrates. Athe_1870, Calhy_0908, Calkr_0826, Calkro_0844, and NA10_0869 from *C. hydrothermalis, C. kristjanssonii, C. kronotskyensis*, and *C. naganoensis*, respectively, where incubated with cellulose (Avicel and filter paper), lignocellulose (switchgrass and poplar), and xylan, along with a no substrate control. Proteins were separated into bound (B) and unbound (U) fractions and visualized with SDS-PAGE; bands here are representative of triplicate trials.

Densitometry analysis of the gels from the tāpirin binding assays supported the conclusions reached from visual inspection (see **Figure 5).** All tāpirins tested had a binding preference for purified cellulose (filter paper more so than Avicel, despite their similar crystallinity (22)), with the exception that NA10_0869 which also bound well to switchgrass. The larger particle size of filter paper (5/16” wide circular disks) compared to Avicel (50 μm particles), as well as filter paper’s higher protein absorption capacity (23, 24), may have been responsible for this difference. Tāpirin binding to lignocellulosic substrates was less than for filter paper and Avicel, likely because of inaccessibility to microcrystalline cellulose in the plant biomasses. As indicated by density of bands on SDS PAGE, more Calhy_0908 bound to Avicel, poplar and filter paper, with the exception being switchgrass **(Figure 4).** Calkr_0826 bound slightly less than Calhy_0908 and more protein bound to switchgrass than for the other tāpirins tested. In fact, the bands of Calhy_0908 and Calkr_0826 are approximately 3 times darker than Athe_1870 and range from 10- to 13-fold more intense than Calkro_0844 and NA10_0869 on filter paper **(Table 3).** Similar trends are noted, but to a lesser extent, on Avicel, with more Calhy_0908 bound than Calkr_0826 and Athe_1870 (2- to 3-fold) and significantly more than NA10_0869 (8-fold) and Calkro_0844 (16-fold) **(Table 3).** It is possible that since the less cellulolytic *Caldicellulosiruptor* species, such *as C. kristjanssonii* and *C. hydrothermalis*, cannot hydrolyze cellulose as well as other species, proximity to these substrates allows these species to exploit the collective lignocellulolytic capacity of *Caldicellulosiruptor* communities in their natural environments.

**Figure 5.**
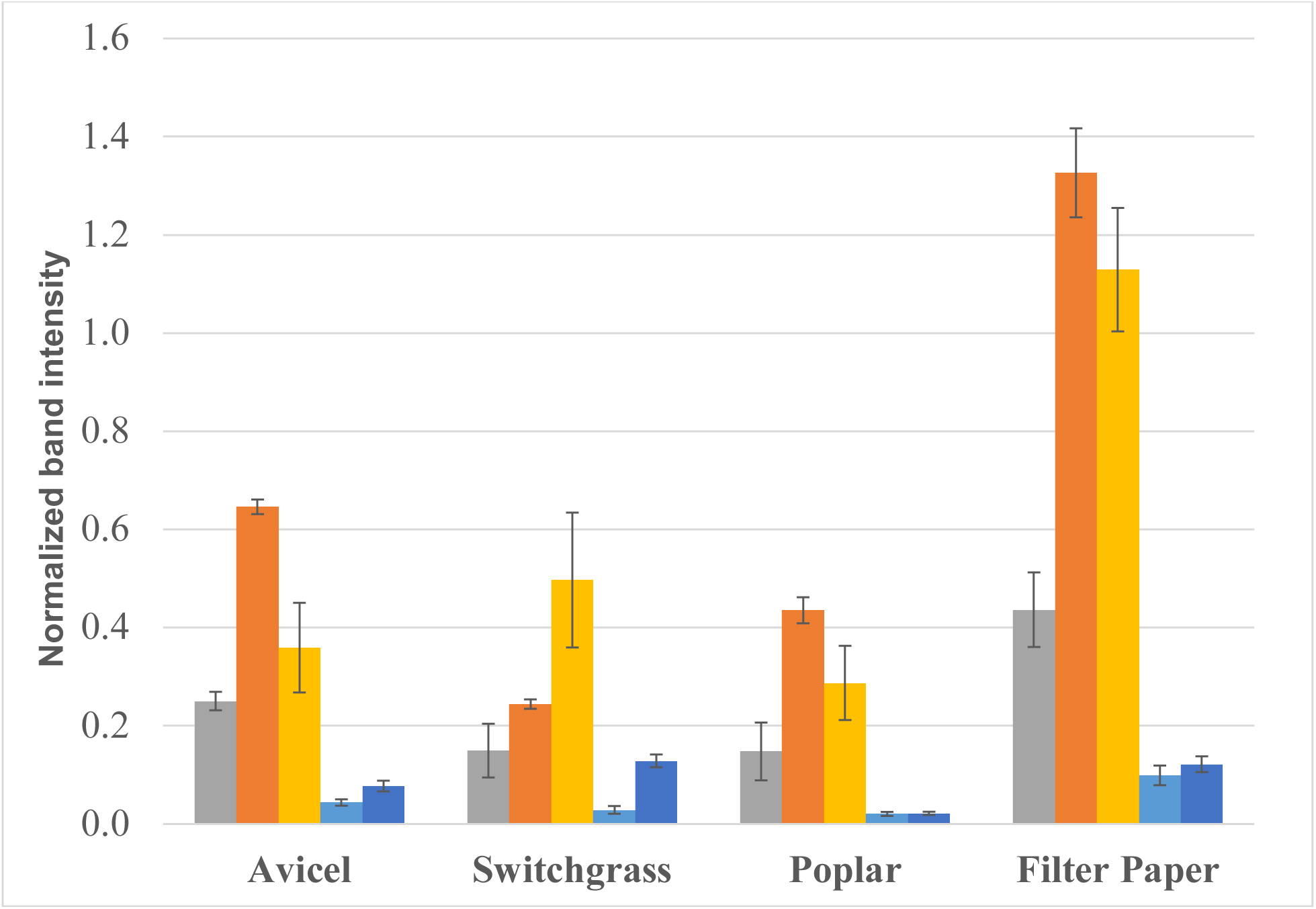
Densitometry of proteins bound to cellulosic substrates as visualized with SDS-PAGE. Bound bands of Athe_1870 (gray), Calhy_0908 (orange), Calkr_0826 (yellow), Calkro_0844 (light blue), and NA10_0869 (dark blue) from *C. bescii, C. hydrothermalis, C. kristjanssonii, C. kronotskyensis*, and *C. naganoensis*, respectively, were quantified and normalized by the 70 kDa Benchmark ladder band on associated SDS-PAGE gel. Error bars represent the standard deviations of the intensities of triplicate samples.

### Structural comparisons of tāpirins from *Caldicellulosiruptor kristjanssonii. Caldicellulosiruptor kronotskyensis*, and *Caldicellulosiruptor hydrothermalis*

The C-terminal domains of tāpirins from *C. kristjanssonii* and *C. hydrothermalis*, Calkr_0826C and Calhy_0908C, respectively, exhibited the same structural architecture as Calkro_0844C from *C. kronotskyensis*, with the core of the domain being a β-helix with a characteristic long loop connecting the ends of the helix **(Figure 6C).** This fold was observed in the Calkro_0844C structure (20) and seems to be a common fold for tāpirins.

**Figure 6.**
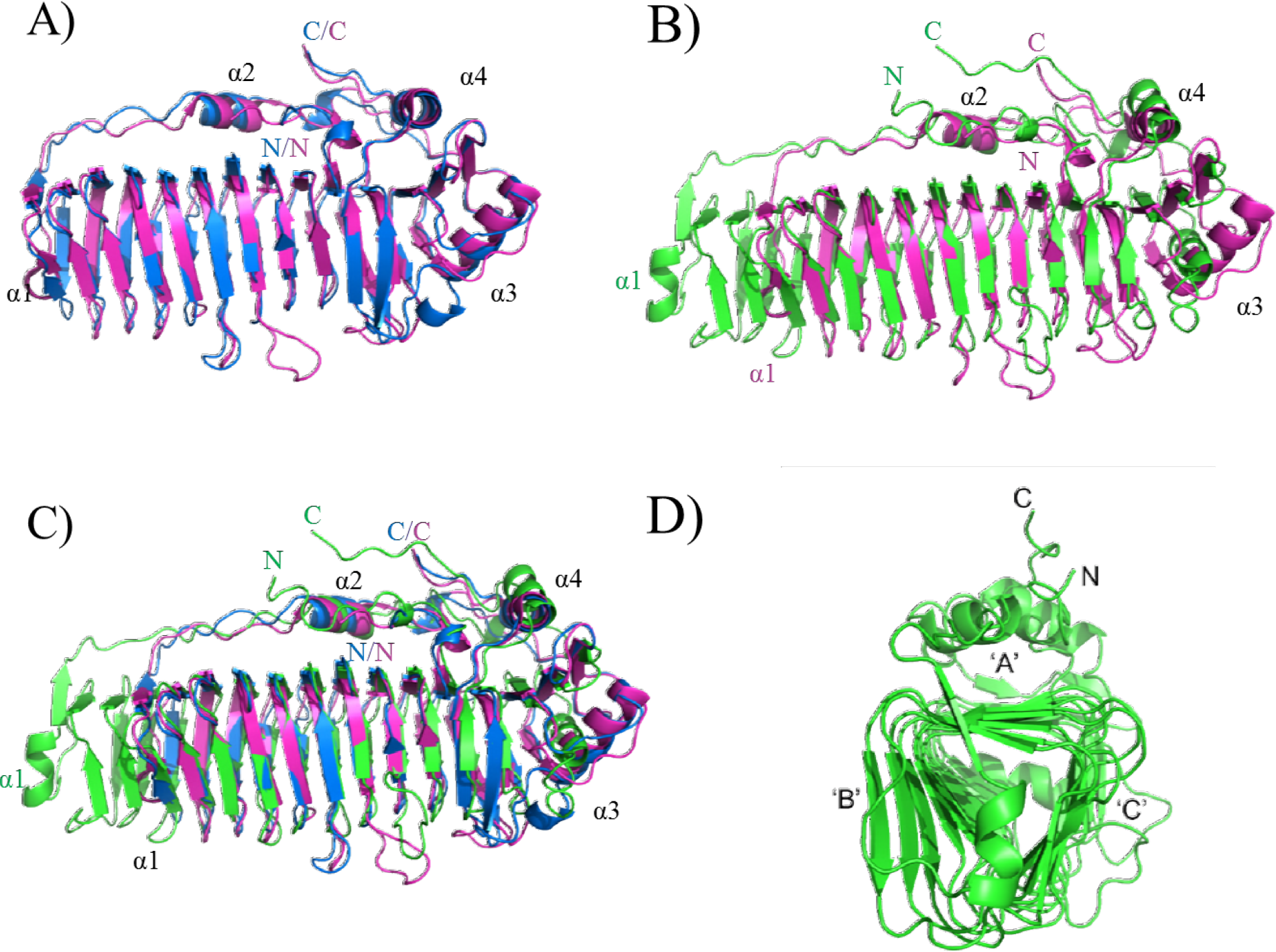
Crystal structures of Calkr_0826, Calhy_0908, and Calkro_0844. A) Calkr_0826 and Calkro_0844 superimposed; B) Calhy_0908 and Calkr_0844 superimposed; C) Calhy_0908, Calkr_0826, and Calkro_0844 superimposed; and D) 90° rotation of Calhy_0908, showing β-helix face designations: ‘A’, ‘B’, and ‘C’. The colors refer to the following tāpirins: green, Calhy_0908 (*C. hydrothermalis*); blue, Calkr_0826 (*C. kristjanssonii*); and magenta, Calkro_0844 (*C. kronotskyensis*). Abbreviations are as follows: α, alpha helix; N, N-terminus; and C, C-terminus. Colors of labels correspond to the colors of the peptides; black font refers to a shared feature across all tāpirins.

The Calkr_0826C structure was superimposed onto Calkro_0844C with the root-mean-square deviation (r.m.s.d.) of 0.854 Å over 1,442 atoms, with both domains having the core ß-helix of the same size **(Figure 6A).** The β-helix in both cases has 11 complete turns, with the longest β-sheet (designated face A, see **Figure 6D)** having 14 β-strands. A few differences, however, are evident: a loop between β8 and β9 in Calkro_0844C is 6 residues longer, β-strand β26 is missing in Calkro_0844C, and the α-helix in Calkro_0844C before β27 is not present in Calkr_0826C. The long loop connecting the ends of the β-helix (β28 to β29, Calkro_0826C) is of the same length (40 residues) in both Calkr_0826C and Calkro_0844C, and features an α-helix located at the same position in the middle. The loop connecting the α2 helix to the β32 strand is 3 amino acid residues shorter in Calkr_0826C, which is compensated by the neighboring loop between β33 and β34, which is 9 residues longer in Calkr_0826C. Also, the loop between β36 and β37 is again 3 residues shorter in Calkr_0826. It is worth noting that the connecting loops of different lengths (β8-β9, α2-β32, β33-β34 and β36-β37) are adjacent and represent about half of the edge between faces B (the next one after A following the direction of the polypeptide, see **Figure 6D)** and C (the face following B, see **Figure 6D)** of the β-helix, opposite to the connecting loop. Similar to the Calkro_0844C, Calkr_0826C has a hydrophobic surface on the face A covered by the connecting loop with multiple flat-on-a-surface aromatic sidechains lined up along the β-sheet **(Figure 6A).**

The C-terminal domain of Calhy_0908C is longer than that of Calkro_0844C and shares the least amount of amino acid sequence homology **(Table 2)** to the other tāpirins examined here. The lack of homology actually led to an incorrect sequence alignment by NCBI-BLAST (25), which in turn translated into an incorrect homology model that was unsuccessfully attempted for molecular replacement. When the molecular replacement attempt failed, the structure of the Calhy_0908C was determined via single-wavelength anomalous diffraction (SAD) using the anomalous signal of iodine atoms incorporated into the crystal after a short soak. Once the structure was determined, a structure-based sequence alignment was a more reliable basis for comparing different tāpirins **(Figure 6B-C).**

Calhy_0908C exhibits the same overall architecture as the other two tāpirins with known structures **(Figure 6C-D):** the β-helix as the core of the domain and a long loop connecting the ends on the β-helix. The major difference between the Calhy_0908C and two other structures is that the β-helix of Calhy_0908C is three turns longer than that of Calkro_0844C and Calkr_0826C, which is made possible by a massive 62 residue single insert in the Calhy_0908C sequence. The rest of the Calhy_0908C (residues 170-378 and 443-578) could be superimposed reasonably well onto Calkro_0844C with the r.m.s.d. of 1.97 Å over 1,020 atoms.

Aside from additional turns of the β-helix, Calhy_0908C has other differences from Calkro_0844C **(Figure 6B).** Similar to the Calkr_0826C vs. Calkro_0844C contrast **(Figure 6A),** the set of loops connecting faces B and C of the β-helix is different. The loop connecting β5 and β6 is 7 residues longer in Calhy_0908C and four more neighboring loops are shorter in Calhy_0908C: the β8-β9 loop, β11-β12 loop, β39-β40 loop, and β42-β43 loop is 7, 2, 2, and 5 residues shorter, respectively. Another difference is the orientation of the helix α3, which while still present in the Calhy_0908C structure, runs almost perpendicular to the corresponding helix of Calkro_0844C. To complement that rearrangement, the connecting loop in Calhy_0908C goes straight to the β36 without an 8-residue ‘detour’ that is present in Calkro_0844C. Also, the β29 strand of Calkro_0844C is not present in Calhy_0908C, with its place taken by the repositioned α3 helix. Similar to both Calkro_0844C and Calkr_0826C, in the hydrophobic face A of Calhy_0908C, the β-helix features a line of aromatic residues **(Figure 7A-C).**

**Figure 7.**
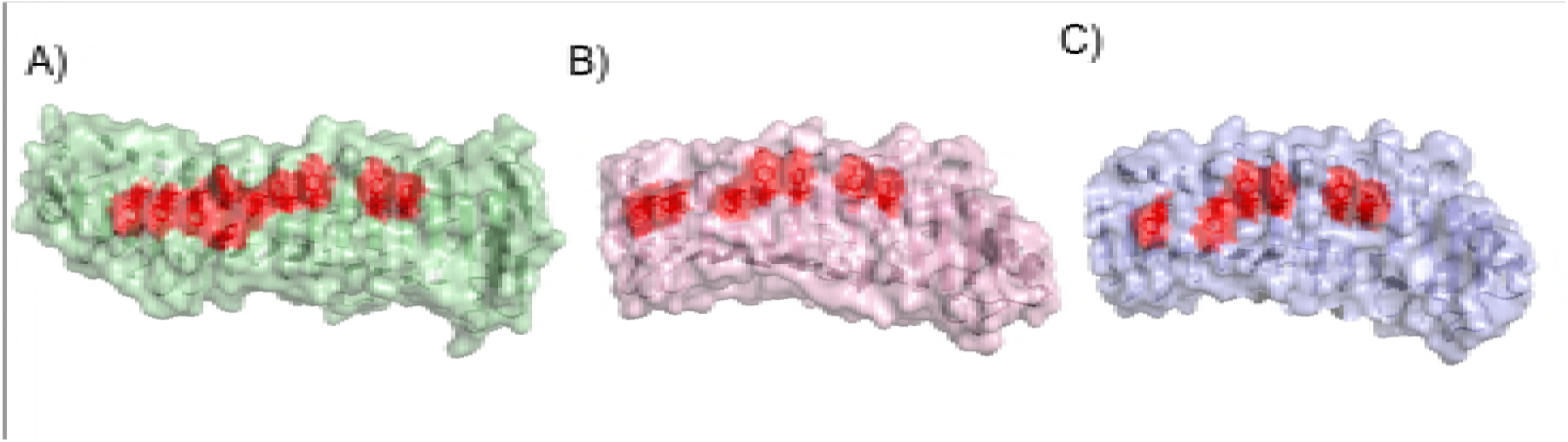
Surface features of tapirins: A) Calhy_0908 (*C. hydrothermalis*),B) Calkro_0844 (*C. kronotskyensis*), and C) Calkr_0826 (*C. kristjanssonii*). Connecting loop and N- and C-termini are removed. Aromatic sidechains are highlighted in red.

### Can the observed differences in tapirin structures explain the differences in cellulose binding

As it was shown in the **Figure 5,** Calhy_0908 binds to more sites on cellulose compared to the other tāpirins examined here. When the structural features of three of these tāpirins are compared **(Figure 6C),** there are two regions where differences are apparent. First, the set of loops, connecting β-strands on the edge between faces B and C of the β-helix, is different in these three proteins, varying in size and chemistry. However, it should be pointed out that Calhy_0908C has the least extensive set of loops, such that most of these are shorter than the corresponding regions in Calkr_0826C and Calkro_0844C. If these loops were responsible for the protein interaction with the cellulose, this difference would leave overall less ‘real estate’ for the possible protein-cellulose interactions. However, that cannot be the case, as Calhy_0908 does indeed seem to bind well to cellulose despite the differences in the loop sets.

Still, another area of interest is the hydrophobic surface found on the face A of the β-helix that is protected by the connecting loop in the crystallized conformations. A possible cellulose-binding mechanism could involve repositioning of the connecting loop upon mechanical contact with the cellulose surface, exposing the line of aromatic sidechains positioned flat on the surface and spaced 5 Å apart, which corresponds to the distance between sugar units in the cellulose chain. This can be seen in **Figure 7,** where Calhy_0908C has the largest hydrophobic surface of the three tāpirins as well as the largest number of aromatic side chains lined up on that surface.

### Localization of tāpirins on the cell surface of *Caldicellulosiruptor*

There is evidence for tāpirin localization on the cell surface, based on fluorescence microscopy using antibodies directed against these proteins **(Figure 8).** Tāpirins are most evident at the poles of the cell, although they also appear to decorate the cellular surface. Given their proposed role as cellulose-binding proteins, especially for initial attachment to substrate, this is consistent with that hypothesis.

**Figure 8.**
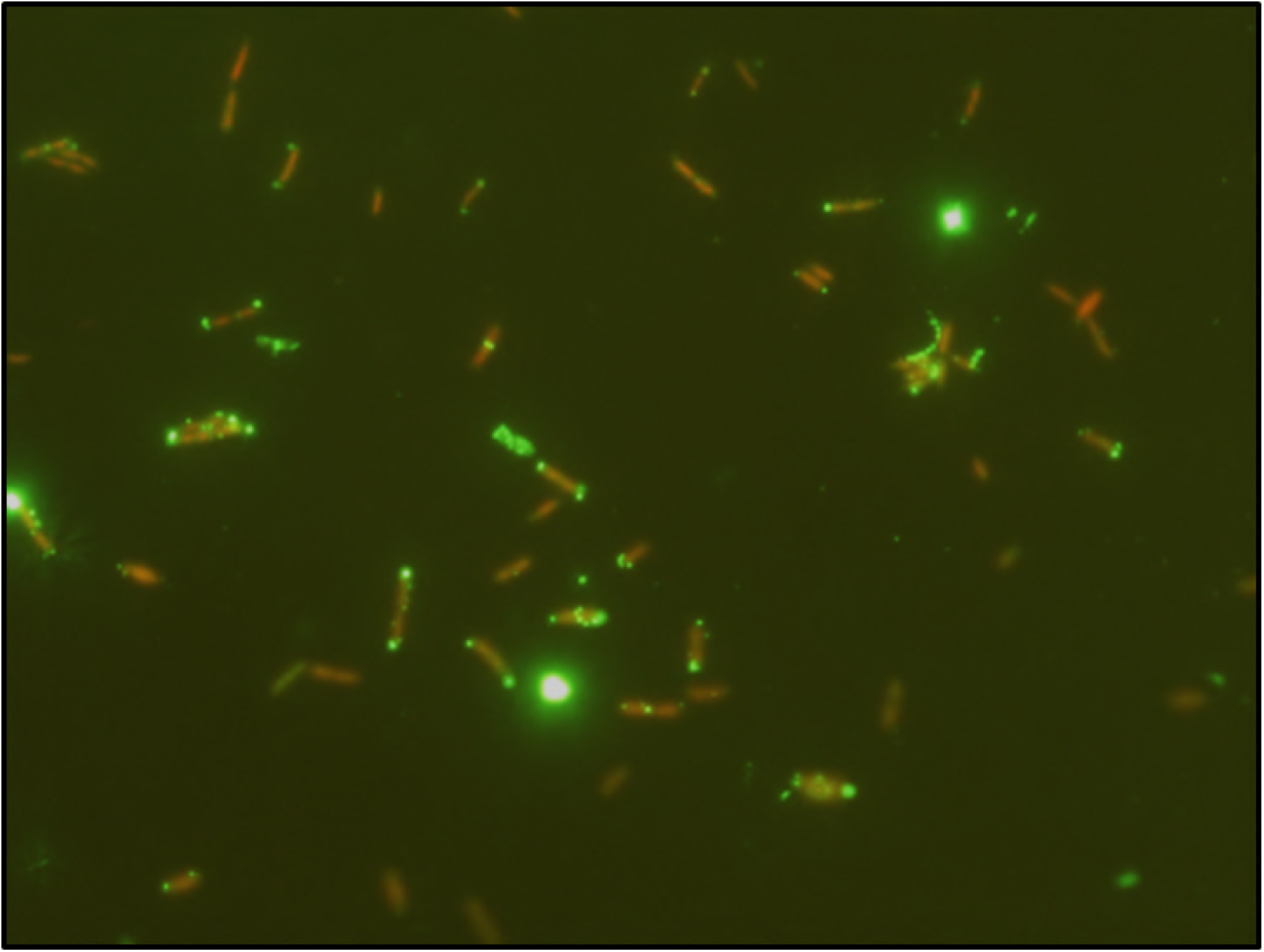
Fluorescence microscopy of *Caldicellulosiruptor kronoyskyensis* using tāpirin antibodies. *C. kronotskyensis* cells (in orange) grown on filter paper were incubated with primary antibody targeting the Calkro_0844 tāpirin, and visualized (in green) with fluorescence microscopy and acridine orange staining (as done in (17)).

### Summary

It interesting that binding assays to cellulosic substrates indicated that Calhy_0908 and Calkr_0826 from the weakly cellulolytic species *C. hydrothermalis* and *C. kristjanssonii*, respectively, appeared to bind better to cellulose than those tāpirins from prolific microcrystalline cellulose degraders. Structures from the C-termini of both Calhy_0908 and Calkr_0826 compared to the previously characterized Calkro_0844 (20) identified clear differences between the tāpirins, including a much longer potential binding platform in Calhy_0908, which contains the largest number of hydrophobic and aromatic residues of the three. This suggests that the less cellulolytic species rely attachment as a key to their survival in their native environments.

While fluorescence microscopy indicates that the presence of tāpirins on the outside of the *Caldicellulosiruptor* cells, their exact cellular location needs to be resolved, especially as this relates to possible association with Type IV pili. Another interesting question going forward is whether tāpirins are a uniquely *Caldicellulosiruptor* feature or whether counterparts are used by other microorganisms to attach to substrates or surfaces.

## Materials and Methods

### Bacterial strains, plasmids, and substrates

The wild type strain of *Caldicellulosiruptor bescii* was obtained from Leibniz Institute DSMZ-German Collection of Microorganisms and Cell Cultures, and *C. bescii* strain MACB1018 and genetic vector, pGL0100, were developed previously (26). *Pyrococcus furiosus* strain COM1 (27) was obtained from Dr. Michael W.W. Adams at the University of Georgia. *Escherichia coli* strains 5-alpha (New England BioLabs) and Rosetta (Millipore Sigma, Merck) were used for plasmid replication and protein expression, respectively. Genes of interest were PCR amplified from extracted genomic DNA, as described previously (28), and Gibson assembly ((29) Gibson assembly master mix, New England BioLabs) or a KLD reaction (KLD Enzyme Mix, New England BioLabs) was used to insert the fragments into plasmids, which had been extracted with ZymoPure midiprep and Zymo Research plasmid miniprep classic kits (Zymo Research). Additionally, Athe_1870 (GenBank™ accession number WP_015908253.1), Calhy_0908 (GenBank™ accession number YP_003992006.1), Calkr_0826 (GenBank™ accession number YP_004025962.1), NA10_0869 (DOE Joint Genome Institute (JGI) Integrated Microbial Genomes & Microbiomes gene ID 2566072544), and Calkro_0844 (GenBank™ accession number YP_004023543.1) genes were *E. coli* codon optimized (without transmembrane domains or signal peptides) with Integrated DNA Technologies (IDT) Codon Optimization Tool (https://www.idtdna.com/CodonOpt), and synthesized by the DOE JGI on a pET_45 plasmid. Sequences of all plasmids and edited portions of final strains were confirmed with Sanger sequencing (Genewiz). Substrates used for growth and binding include: Avicel PH-101 (FMC BioPolymer), Cave-in-Rock switchgrass *(Panicum virgatum* L. from fields in Monroe County, IA, retrieved by the National Renewable Energy Laboratory, and ground and sieved using a Wiley mill (Thomas Scientific) and 40/80 mesh, respectively), beechwood xylan (Sigma-Aldrich), and poplar *(Populus trichocarpa*, obtained from Vincent Chiang (30))

### Expression of tapirin proteins in *E. coli*

Expression plasmids with synthesized Athe_1870, Calhy_0908, Calkr_0826, NA10_0869, and Calkro_0844 genes were transformed into *E. coli* (strain 5-alpha with 50 μg/ml carbenicillin section and strain Rosetta with both 50 μg/ml carbenicillin and 34 μg/ml chloramphenicol selection), and cultured on Luria-Bertani (LB – 10 g/liter sodium chloride, 10 g/liter tryptone, and 5 g/L yeast extract) liquid medium or agar (1.5% wt/vol) plates at 37°C. For the production of protein, the cultures were grown in ZYM-5052 autoinduction medium (31) with chloramphenicol and carbenicillin in up to 1-3L volumes at 37°C 250 RPM for 18-24 hours and harvested with centrifugation of 6000 x g for 10 minutes. Cells were then re-suspended in 100 mL of 20 mM sodium phosphate, pH 7.4, 0.5 mM sodium chloride, and 5 mM imidazole, lysed with a French Press at 16,000 psig, heat-treated at 65°C for 30 minutes, and centrifuged at 25,000 x g for 30 minutes. Protein in soluble fraction was purified with 5-ml HisTrap HP nickel-Sepharose immobilized metal affinity chromatography column (GE Healthcare operated according to the manufacturer’s instructions) using a Biologic DuoFlow FPLC (Bio-Rad), and then stored at 4°C. Protein concentration was determined by the Bradford assay (32) and protein purity was visualized along with a Benchmark protein ladder (Life Technologies) by SDS-PAGE using 4-15% Mini-PROTEAN^®^ TGX Stain-Free™ Precast Gels (Bio-Rad).

### Expression and purification of C-terminal tāpirins, Calkr_0826C and Calhy_0908C

Purified protein was buffer-exchanged into 0.4 mM calcium chloride, 0.15 M sodium chloride, and 50 mM tris-chloride (pH 8.0) reaction buffer and treated with thermolysin (1 mg/ml in same buffer Promega), as described in (20). A thermolysin-to-protein ratio of 1:500 and treatment times of 5 to 30 minutes at 70°C were used to effectively lyse protein to produce the C-terminal portion of the tāpirin (i.e. Calkr_0826C for *C. kristjanssonii* and Calhy_0809C for *C. hydrothermalis).* Reactions were halted by storing samples on ice and then long term at 4°C. Samples were imaged with SDS-PAGE as described above to verify cleavage. For the crystallization of Calkr_0826C and Calhy_0908C, cleaved products were further purified using an ÄKTA protein purification system (GE Healthcare Life Sciences) and Superdex 75 pg (16/60) size exclusion chromatography column in 20 mM Tris pH 7.5 and 100 mM sodium chloride.

### Crystallization

The crystals of Calkr_0826C and Calhy_0908C were initially obtained with sitting drop vapor diffusion using a 96-well plate with PEG ion HT screen from Hampton Research (Aliso Viejo, CA). 50 μL of well solution was added to the reservoir, and drops were made with 0.2 μL of well solution and 0.2 μL of protein solution using a Phoenix crystallization robot (Art Robbins Instruments, Sunnyvale, CA). The Calkr_0826C crystals were grown at 20°C using an optimization screen containing 0.1 M citric acid pH 3.0 to 4.0 and 10% to 15% w/v polyethylene glycol (PEG) 3350 (best crystals appeared in pH range from 3.1 to 3.2 and PEG 3350 14-15%). The protein solutions contained 12 mg/mL of protein, 20 mM Tris pH 7.5, 100 mM sodium chloride, 2% of the Hampton Research Tacsimate pH 4 mix and 5 mM of each of zinc acetate, potassium chloride, magnesium chloride, and calcium chloride. The Calhy_0908C crystals were grown at 20°C using an optimization screen containing 5 mM – 35 mM zinc acetate and 15% to 24% w/v PEG 3350 (best crystals appeared in 0.015 M zinc acetate and PEG 3350 concentration of 17-18%). The protein solutions contained 7.5 mg/mL of protein, 20 mM Tris pH 7.5, 100 mM NaCl and 2% of the Hampton Research Tacsimate pH 7 mix.

All crystals were soaked in well solution with PEG 3350 increased to 25% along with 5-10% ethylene glycol added for the cryo protection. For the purpose of structure determination, an iodine derivative was obtained for the Calhy_0908C by quick soaking the crystals in the cryoprotectant described above with 0.1 M potassium iodide added.

### Crystallography data collection and processing

The crystals of Calkr_0826C and Calhy_0908C were flash frozen in a nitrogen gas stream at 100 K before home source data collection using an in-house Bruker X8 MicroStar X-Ray generator with Helios mirrors and Bruker Platinum 135 CCD detector. Data were indexed and processed with the Bruker Suite of programs version 2014.9 (Bruker AXS, Madison, WI).

### Crystal structure solution and refinement

Intensities were converted into structure factors and ‘free’ sets of the reflections (5% of the reflections for Calkr_0826C and 2% for Calhy_0908C) were flagged for R_free_ calculations using programs F2MTZ, Truncate, CAD and Unique from the CCP4 package of programs (33). The structure of the Calkr_0826C was solved by MOLREP (34) using Calkro_0844_C (20) (PDB ID 4WA0) as a search model. Crank2 (35) was used to solve the structure of the Calhy_0908C utilizing iodine single-wavelength anomalous dispersion (36). Refinement and manual correction was performed using REFMAC5 (37) version 5.8.158, PHENIX (38) version 1.11 and Coot (39) version 0.8.8. The MOLPROBITY method (40) was used to analyze the Ramachandran plot, and root mean square deviations (rmsd) of bond lengths and angles were calculated from ideal values of Engh and Huber stereo chemical parameters (41). Wilson B-factor was calculated using CTRUNCATE version 1.15.10 (33). The data collection and refinement statistics are shown in Table **4.4.**

### Tāpirin binding assays

Recombinant proteins, both full length and truncated, were tested for attachment to various substrates in triplicate, as described in (20). All substrates were initially soaked with 100 mL of 50 mM MES and 3.9 mM sodium chloride at pH 7.2 (‘binding buffer’) overnight, and then subsequently dried overnight (both steps at 70°C). Nine mg of washed substrates were mixed with 40 μg of purified tāpirin protein and incubated in a thermomixer (Eppendorf) at 70°C and 500 rpm for one hour. Samples were then centrifuged 13000 x g and separated into ‘unbound’ (supernatant) and ‘bound’ (pellet) fractions. The bound fraction was then washed (re-suspending the substrate in binding buffer with vortexing, centrifuging mixture at 13000 x g, and discarding the supernatant) four times before being finally re-suspended in 250 μL of buffer. Equal volumes of bound or unbound sample were mixed with 2x Laemmli sample buffer and 5% 2-mercaptoethanol, and boiled for 30 minutes. Samples were then loaded on a SDS-PAGE gel as described above. Densitometry was completed by using ImageJ (42) to analyze band intensity (keeping contrast across gels consist) and by normalizing all bands to the 70 kDa Benchmark protein ladder (Life Technologies) band present on each gel.

### Deletion of tāpirin and/or pili genes in *C. bescii*

Knockout (KO) vectors were constructed with Gibson Assembly with pGL100 (26) as the backbone with *pyrE* (Athe_1382). Flanking regions outside Athe_1870-1871, and Athe_1870-1885 were PCR-amplified using *C. bescii* MACB1018 genomic DNA as a template, while the vector backbone and kanamycin resistance gene (HTK) and SLP promoter (Athe_2303) and were PCR-amplified from a template plasmid before using Gibson Assembly to assemble the knock out vectors (see below). A new genetic vector, pLLL023, was also generated using a KLD reaction from pGL100 to create a plasmid without P_slp_-HTK on the backbone, such that P_slp_-HTK could be inserted into the genome. Primers used for vector construction and genetic screening are included in **Table 4.5.** The resulting vectors were generated for the following strains: pLLL024 for RKCB136 (Athe_1870-1871 KO and P_slp_-HTK knock-in (KI)), and pLLL012 for RKCB135 (Athe_1870-1885 KO). After construction, plasmids were methylated with purified recombinant M.CbeI (as described previously in (26, 43)).

*C. bescii* genetic strains were all cultured anaerobically in either low osmolality defined (LOD) or complex (LOC) medium (44) with cellobiose in liquid cultures at 75°C with a nitrogen headspace and on 1.5% wt/vol agar plates in an anaerobic chamber (Coy Laboratory) at 65°C. To transform KO plasmids into *C. bescii* strain MACB1018 (26), 1 μg of plasmid DNA was added to 50 μl aliquots of competent cells (prepped as described previously (26) in LOD media supplemented with 1 × 19 amino acid solution (27)) at room temperature. Cells were then electroporated with a Gene Pulser II system with a Pulse Controller Plus module (Bio-Rad) in 1-mm-gap cuvettes (USA Scientific) at 25 μF, 200 Ω, and 2 kV before being passaged to 10 mL of preheated LOC medium for recovery at 75°C. After one hour, all recovery media was passaged to selective media (LOC with 50 μg/mL kanamcyin) and incubated until growth was noted (typically 1 to 3 days) at 75°C. Growing transformants were screened with PCR and then passaged multiple times into liquid selective medium and/or plated on selective media for purification. Second crossovers were then selected by plating transformed cells onto LOD medium containing 4 mM 5-fluoroorotic acid (5-FOA), 50 μg/mL kanamycin (for HTK KI strains), and 40 mM uracil. Successful PCR-screened colonies were plate purified on the same second-crossover plating media without 5-FOA.

### Whole cell binding assay

*Caldicellulosiruptor* cells were cultured at 75°C in 100 mL of modified 671 medium with 5 g/L Avicel, as described previously. Cultures were measured for cell density with acridine orange epifluorescence microscopy, as described previously (45), before the entire sample was initially centrifuged at 400 x g for 5 minutes to lightly pellet Avicel. The supernatant was removed and cells were harvested by centrifugation at 6000 x g for 10 minutes before being concentrated to 5×10^8^ to 1×10^9^ cells/ml in 671d medium. 1 mL of cells were added to 10 mg of Avicel for 1 hour in a thermomixer (Eppendorf) at 70°C; a control was also completed without any substrate (only media and cells) and all samples were completed in triplicate. The supernatant counting unbound cells were then separated and enumerated again with acridine orange epifluorescence microscopy, with the average planktonic cell densities were compared using a t-test.

## Acknowledgements

This work was supported by the BioEnergy Science Center (BESC), a U.S. Department of Energy Bioenergy Research Center supported by the Office of Biological and Environmental Research in the DOE Office of Science. LL Lee acknowledges support from a National Science Foundation Graduate Research Fellowship and a NIH T32 Biotechnology Traineeship (GM008776-11). Synthetic genes for the expression of the tāpirins were provided by the U.S. Department of Energy Joint Genome Institute, a DOE Office of Science User Facility, which is supported by the Office of Science of the U.S. Department of Energy under Contract No. DE-AC02-05CH11231.

